# Alanine dependence of trans-translation contributes to riboregulation of mycobacterial antibiotic recalcitrance genes

**DOI:** 10.1101/2025.10.10.681568

**Authors:** Andrea Majstorovic, Helene Botella, Sarah M. Schrader, Yaroslav Lavrynchuk, Xinguo Chen, Enkai Jin, Jason Taslim, Rudy Antoine, Alain Baulard, Christophe Guilhot, Przemysław Płociński, Sandra L. Wolin, Julien Vaubourgeix

## Abstract

Antibiotic recalcitrance refers to a slower rate of death for either a bacterial population or a subpopulation of cells upon antibiotic exposure. It complicates treatment of many bacterial infections by contributing to treatment length, treatment failure, disease recurrence, and the emergence of antimicrobial resistance (AMR). Thus, blocking antibiotic recalcitrance could be a powerful strategy for improving treatment outcomes and reducing AMR rates. Here, using a forward genetic method for the isolation of antibiotic-recalcitrant mutants, we isolated two *Mycobacterium smegmatis* strains with mutations in the tRNA-modifying enzyme adenine-N(1)-methyltransferase. Both mutants were recalcitrant to proteostasis-perturbing antibiotics. We linked these phenotypes to upregulation of the transcriptional regulator WhiB7, highlighting its role as a point of convergence in the regulation of multiple mechanisms of antibiotic recalcitrance and resistance. Further, we identified a mechanism by which the amino acid alanine couples *trans*-translation to ribosome regulation-dependent, WhiB7-mediated expression of antibiotic resistance and recalcitrance genes, allowing bacterial cells to engage seemingly mutually exclusive mechanisms of survival upon exposure to stress.

## INTRODUCTION

Antibiotics offer a cure for many infectious diseases and enable modern medical interventions. However, antimicrobial resistance (AMR), which directly caused 1.3 million deaths in 2019 with even grimmer projections, threatens to reverse these gains (Murray *et al*., 2022; Ho *et al*., 2025). In September 2024, the United Nations (UN) General Assembly convened a High-Level Meeting on AMR—the second in history—to reaffirm the threat AMR represents to global health and food security and press UN members to execute plans to combat it.

AMR refers to the growth of bacteria in the presence of an antibiotic due to acquisition of a stably heritable mutation or new genetic material. Antibiotic tolerance and persistence—collectively referred to as high survival (Schrader *et al*., 2021) or antibiotic recalcitrance (Fisher, Gollan and Helaine, 2017)—are phenomena that are related to AMR but describe increased survival upon exposure to an antibiotic for either the entire bacterial population (tolerance) or a subpopulation of bacterial cells (persistence) (Balaban *et al*., 2019; Schrader, Vaubourgeix and Nathan, 2020; Schrader, Botella and Vaubourgeix, 2023) (Fig. 1A). Both complicate treatment of bacterial infections by prolonging treatment and causing treatment failure and disease recurrence (Etthel M. Windels *et al*., 2019; Huemer *et al*., 2020). Furthermore, prolonged survival under antibiotic pressure along with increased mutation rates in such conditions facilitate the development of AMR in tolerant populations and persisters (Levin-Reisman *et al*., 2017; Sebastian *et al*., 2017; Etthel Martha Windels *et al*., 2019; Liu *et al*., 2020; Russo *et al*., 2022; Wilmaerts *et al*., 2022). Therefore, therapeutic strategies that target tolerance and persistence could improve treatment outcomes in infectious disease and curb AMR.

**Figure 1.**
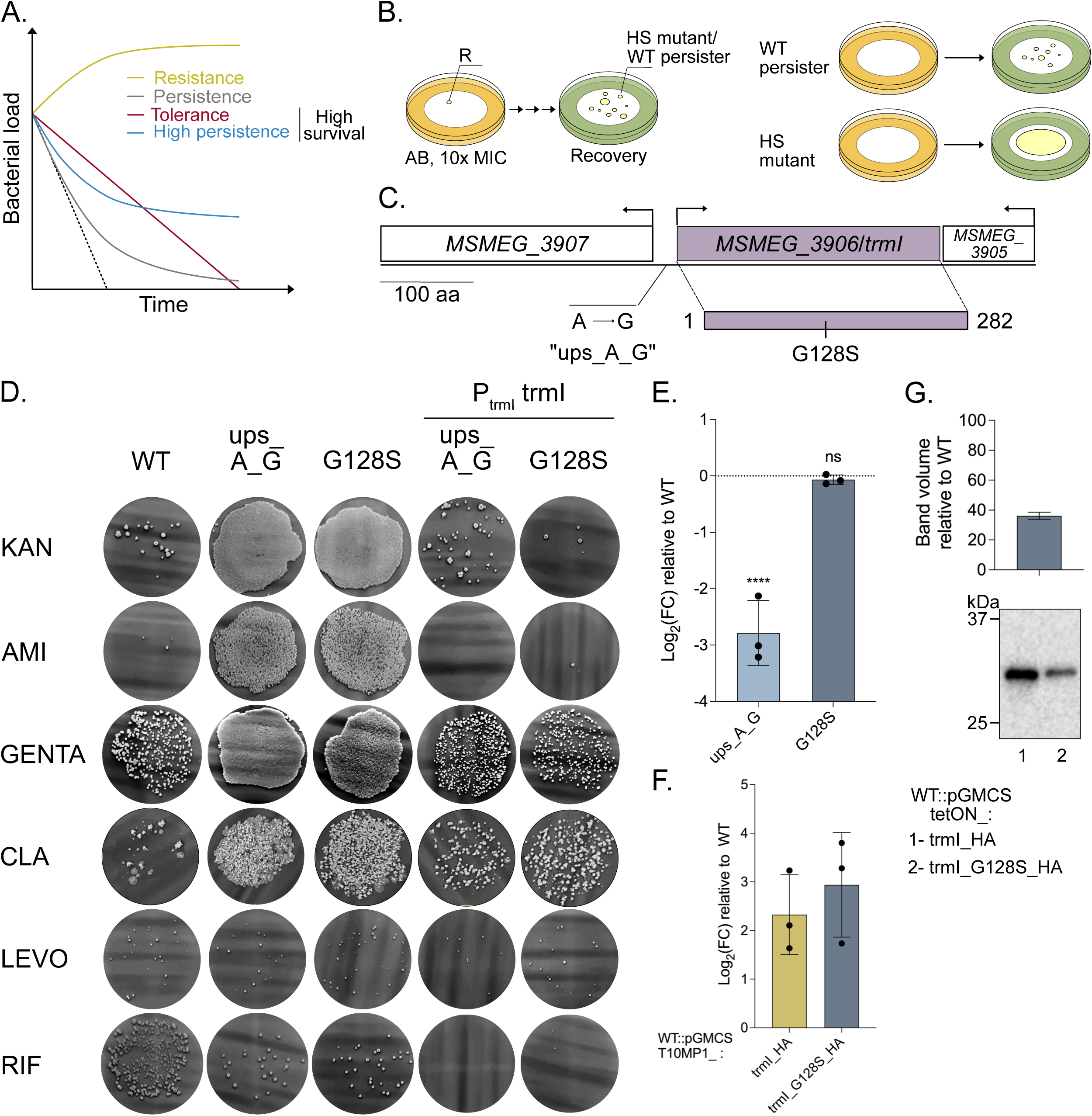
Loss-of-function mutations in a gene encoding a tRNA methyltransferase lead to high survival upon exposure to antibiotics. **A.** Manifestations of antimicrobial resistance/recalcitrance. Graphs represent bacterial load over time in the presence of an antibiotic for resistant and tolerant populations and populations containing persisters at basal (persistence) or increased (high persistence) levels. **B.** Schematic representation of the screening workflow initially published in (Schrader *et al*., 2021) that was used for selection and identification of the adenine-N(1)-methyltransferase mutants. R, genetically resistant mutant; HS, high-survival mutant; WT, wild-type; AB, antibiotic. **C.** Schematic representation of the *MSMEG_3906*/*trmI* locus in *Msm*. The location and nature of the mutations found in the two isolated adenine-N(1)-methyltransferase mutants are indicated; one mutant (ups_A_G) has an A-to-G mutation 15 bp upstream of *MSMEG_3906*, while the other (G128S) has a C-to-T mutation leading to a glycine-to serine substitution at position 128 within the coding sequence. **D.** Survival of WT *Msm* (WT), the adenine-N(1)-methyltransferase mutants (ups_A_G and G128S), and their respective complemented strains (ups_A_G::P_­­­­_*ms_trmI* and G128S::P_­­­­_*ms_trmI*) upon exposure to various antibiotics (KAN, kanamycin; AMI, amikacin; GENTA, gentamycin; LEVO, levofloxacin; RIF, rifampicin; CLA, clarithromycin). **E.** Expression level of *MSMEG_3906* (*trmI*) in the adenine-N(1)-methyltransferase mutants compared to WT *Msm*. The dotted line represents no change relative to the WT transcript level. FC, fold change. ****, p < 0.0001. ns, not significant. **F.** and **G.** Expression level of *MSMEG_3906* (*trmI*) transcript (F) and the trmI protein (G) in WT *Msm* bearing an allele for either HA-tagged, WT Trml or HA-tagged, G128S-mutated TrmI. The bar graph in G represents the surface area of the Trml_HA band seen in the Western blot shown beneath the bar graph divided by the surface area of the Trml_G128S_HA band; equal amounts of protein extracts were loaded for each strain. This is quantified in the right panel. In (G), error bars represent +/− SD, means of three replicates, one of which is shown in (F).

Tolerance and high persistence—which refers to an increased proportion of persisters in an otherwise antibiotic-susceptible population—can each result from acquisition of heritable genetic changes (mutations or stably retained genes) or conditional induction of diverse, incompletely understood mechanisms in bacteria that are genetically identical to those that are susceptible. Of note, a genetic change that causes high persistence is carried by all bacteria in a population and serves to increase the chance that an individual cell will enter a phenotypic state that conveys persistence. The systematic isolation of tolerant and highly persistent mutants, which do not grow at a population level during antibiotic exposure, is challenging because they are difficult to separate from resistant mutants, which multiply during antibiotic exposure. We previously developed a four-step, *in vitro*, forward genetic method called Antibiotic Survivor Separation In Space and Time (ASSIST) that allows for spatiotemporal separation of resisters from survivors without an increase in the minimal inhibitory concentration (MIC), which include tolerant and highly persistent mutants (Schrader *et al*., 2021). This forward-genetic method can be applied to any organism cultivable on solid medium and permits isolation and subsequent validation of mutants that display tolerance or high persistence (high-survival mutants). The method involves seeding filters with bacteria and placing them onto agar plates containing 10x the MIC of an antibiotic for a period pre-determined to kill > 99% of a wild-type (WT) population. The filters are then transferred to agar plates containing 2x the MIC of the same antibiotic. During these exposure periods, high-level (ΔMIC > 10) and then low-level (2 > ΔMIC < 10) resisters are able to grow and form visible colonies. Next, filters are transferred to growth-permissive agar containing 0.4% charcoal for 24 hours to remove residual antibiotic before final transfer to growth-permissive agar for recovery of tolerant and persistent bacteria. Colonies that emerge during the recovery period are picked and outgrown in liquid medium as candidate high-survival mutants. To distinguish between WT persisters and high-survival mutants, each candidate without an increase in MIC for the drug used in selection is seeded onto a filter, and the filter transfers are repeated with omission of the 2x MIC exposure step. Candidates with a reproducible > 10-fold increase in the number of survivors compared to a WT control qualify as high-survival mutants (Fig. 1B).

Using ASSIST, we previously isolated high-survival mutants in *Mycobacterium smegmatis* (*Msm*) in which perturbation of the arginine biosynthesis pathway generated three forms of resistance or recalcitrance upon exposure to diverse antibiotics, each mediated by induction of the transcriptional regulator WhiB7: high persistence and tolerance to kanamycin, high survival upon exposure to rifampicin, and resistance to macrolides. Although the mechanisms for all these forms of resistance/recalcitrance converged on WhiB7, the direct mediators within the WhiB7 regulon differed for each antibiotic. We showed that the aminoglycoside-modifying enzyme Eis mediated high persistence and tolerance to kanamycin, while high survival upon exposure to rifampicin and resistance to macrolides were likely mediated by efflux pumps. These results, together with work by others (Bernard *et al*., 2024), established WhiB7 as a mediator of antibiotic tolerance and persistence, adding to its known role in resistance to some antibiotics (Morris *et al*., 2005; Burian *et al*., 2012, 2013; Hurst-Hess, Rudra and Ghosh, 2017; Schildkraut *et al*., 2022).

Here, we studied two high-survival *Mycobacterium smegmatis* strains that we had previously isolated (Schrader *et al*., 2021) with mutations in the tRNA-modifying enzyme adenine-N(1)-methyltransferase. Both mutants were tolerant to aminoglycosides and macrolides. Similar to our previously published mutants with defects in arginine biosynthesis (Schrader *et al*., 2021), we showed that these phenotypes were mediated by upregulation of the transcriptional regulator WhiB7. This finding positions WhiB7 as a point of convergence for multiple mechanisms of antibiotic recalcitrance and resistance (Schrader, Botella and Vaubourgeix, 2023). Further, we identified a novel mechanism by which the amino acid alanine couples *trans*-translation to ribosome regulation-dependent, WhiB7-mediated expression of antibiotic resistance and recalcitrance genes, allowing mycobacteria to engage seemingly mutually exclusive mechanisms of survival upon exposure to stress.

## RESULTS

### Loss-of-function mutations in a gene encoding a tRNA methyltransferase result in high survival to aminoglycosides

Using ASSIST, we previously isolated two mutants that had mutations in or near the gene *MSMEG_3906*, which encodes a tRNA methyltransferase whose homologs in other microbial species have been named TrmI. We thus refer to *MSMEG_3906* (encoding MSMEG_3906) as *ms_TrmI* (ms_TrmI) hereafter. One mutant (ups_A_G) had an A-to-G mutation 15 bp upstream of *ms_trmI*, while the other (G128S) had a C-to-T mutation leading to a glycine-to serine substitution at position 128 within the coding sequence (Fig. 1C). Both mutants were selected using kanamycin, an aminoglycoside that binds to the 30S ribosomal subunit and disrupts protein synthesis. In addition, both mutants displayed high survival upon exposure to the aminoglycosides amikacin and gentamicin and to the macrolide clarithromycin, which belongs to another class of protein synthesis inhibitors that bind to the 23S rRNA of the 50S ribosomal subunit. The mutants did not display high survival upon exposure to the DNA gyrase inhibitor levofloxacin or the RNA polymerase inhibitor rifampicin (Fig. 1D). We confirmed that the minimum inhibitory concentrations (MICs) of kanamycin, amikacin, gentamicin, clarithromycin, levofloxacin and rifampicin were unchanged in the mutants (Supplemental Fig 1A). Addition of a WT allele of *ms_trmI* under the control of its native promoter reduced survival upon exposure to aminoglycosides and macrolides back to WT levels (Fig. 1D). Next, we sought to determine the impact of each mutation on *ms_trmI*. Using qRT-PCR, we showed that the A-to-G substitution mutation 15 bp upstream of *ms_trmI* led to an approximately 85% decrease in its expression (Fig. 1E). In contrast, the mutation that led to a G-to-S amino acid substitution at position 128 of the coding sequence did not affect the expression of *ms_trmI*. Protein structure modelling analysis of ms_TrmI based on the solved structure of the *Mycobacterium tuberculosis* (*Mtb*) homolog did not support a role of the glycine at position 128 in homotetramer formation or binding of the S-adenosyl-methionine cofactor, both of which are required for the protein to carry out its tRNA-modifying function (Gupta *et al*., 2001; Varshney *et al*., 2004). To explore whether the G-to-S substitution mutation had an effect on ms_TrmI protein abundance, we expressed an HA-tagged version of the WT or mutated ms_TrmI protein in WT *Msm* under control of the inducer anhydrotetracycline (atc) (Supplemental Fig. 1B). Although expression of either allele led to similar mRNA transcript levels, the abundance of the ms_TrmI_G128S protein was only 36% that of WT MSMEG_3906 when assessed 12 hours after removal of atc (Fig. 1F and 1G), suggesting that ms_TrmI_G128S exhibits decreased stability and/or more rapid degradation. These results indicated that both mutations led to decreased levels of ms_TrmI.

Having established that loss-of-function mutations in *ms_trmI* led to high survival upon exposure to protein synthesis inhibitors, we next sought to understand the molecular consequences of impaired TrmI function with regard to its role in antibiotic tolerance. Given the similar effect of the two *ms_trml* mutations on MSMEG_3906 abundance, we performed all subsequent experiments using the ups_A_G mutant.

### Loss of function of ms_TrmI leads to hypomodification of tRNAs, thereby impacting tRNA stability

In *Thermus thermophilus*, loss of *trmI* results in a growth defect at 80 degrees Celsius (Droogmans *et al*., 2003), and TrmI maintains stability of initiator tRNAs in *Saccharomyces cerevisiae* (Anderson *et al*., 1998; Kadaba *et al*., 2004). In addition, previous work showed that *rv2118c*, the homolog of *ms_TrmI* in *Mtb*, is a tRNA (adenine(58)-N(1))-methyltransferase that methylates initiator tRNA when expressed in *Escherichia coli* (Varshney *et al*., 2004). Given this, we evaluated tRNA methylation by ms_TrmI in WT *Msm* and the ups_A_G mutant. The adenine at position 58 of the TλφϑC loop (A58) is conserved on all tRNAs in *Msm* (https://trna.ucsc.edu/tRNAviz/summary/#) (Fig. 2A). Using small RNA sequencing, we measured the rate of base misreading at position 58 during reverse transcription into cDNA across all tRNAs as a proxy for epitranscriptional modification abundance: presence of an epitranscriptional modification at position 58 prevents accurate recognition of the base as adenine, resulting in a misread, while non-modified A58 is correctly recognized (Ebhardt *et al*., 2009; Helm *et al*., 2021; Lucas *et al*., 2024) (Fig. 2B, schematics). Decreased expression levels of *ms*_*trmI* in the ups_A_G mutant led to decreased levels of m^1^A58 methylation of most but not all tRNAs (mean absolute deviation WT vs ups_A_G: 34.5), although expression of a WT copy of *ms_trmI* under the control of its native promoter only partially restored WT levels of tRNA modification at position A58 (mean absolute deviation WT vs ups_A_G_*trmI*: 19.2) (Fig. 2B; Supplemental Fig. 2A). These results also demonstrated that unlike for the *Mtb* homolog upon expression in *E. coli*, m^1^A58 tRNA methylation by ms_TrmI was not restricted to initiator tRNAs (Varshney *et al*., 2004). Next, we aimed to determine whether lack of tRNA methylation at position 58 led to tRNA instability given the reported role of *S. cerevisiae* TrmI in maintaining initiator tRNA stability. To test this, we extracted tRNAs from WT *Msm*, the ups_A_G mutant, and the complemented strain under acidic conditions to maintain aminoacylation. After fractionation in acid polyacrylamide gels under denaturating conditions, overall levels of three tRNAs—tRNA^Arg(UCU)^, tRNA^His(GUG)^, and tRNA^Ala(CGC)^ that our small RNAseq data showed were modified by ms_TrmI—were decreased in the ups_A_G mutant, suggesting decreased stability of these tRNAs. In addition, decreased tRNA methylation in the ups_A_G mutant led to a shift in migration of both charged (aminoacylated) and uncharged tRNA that was complemented by addition of a WT copy of *ms_trmI* under control of its native promoter (Fig. 2C). The ratio of charged-to-uncharged tRNAs was unaffected by decreased methylation in the ups_A_G mutant, suggesting no difference in aminoacylation. Consistent with our small RNAseq data showing lack of modification of tRNA Val (CAC) by ms_TrmI, we did not observe a difference in overall level or a shift in migration for this tRNA.

**Figure 2.**
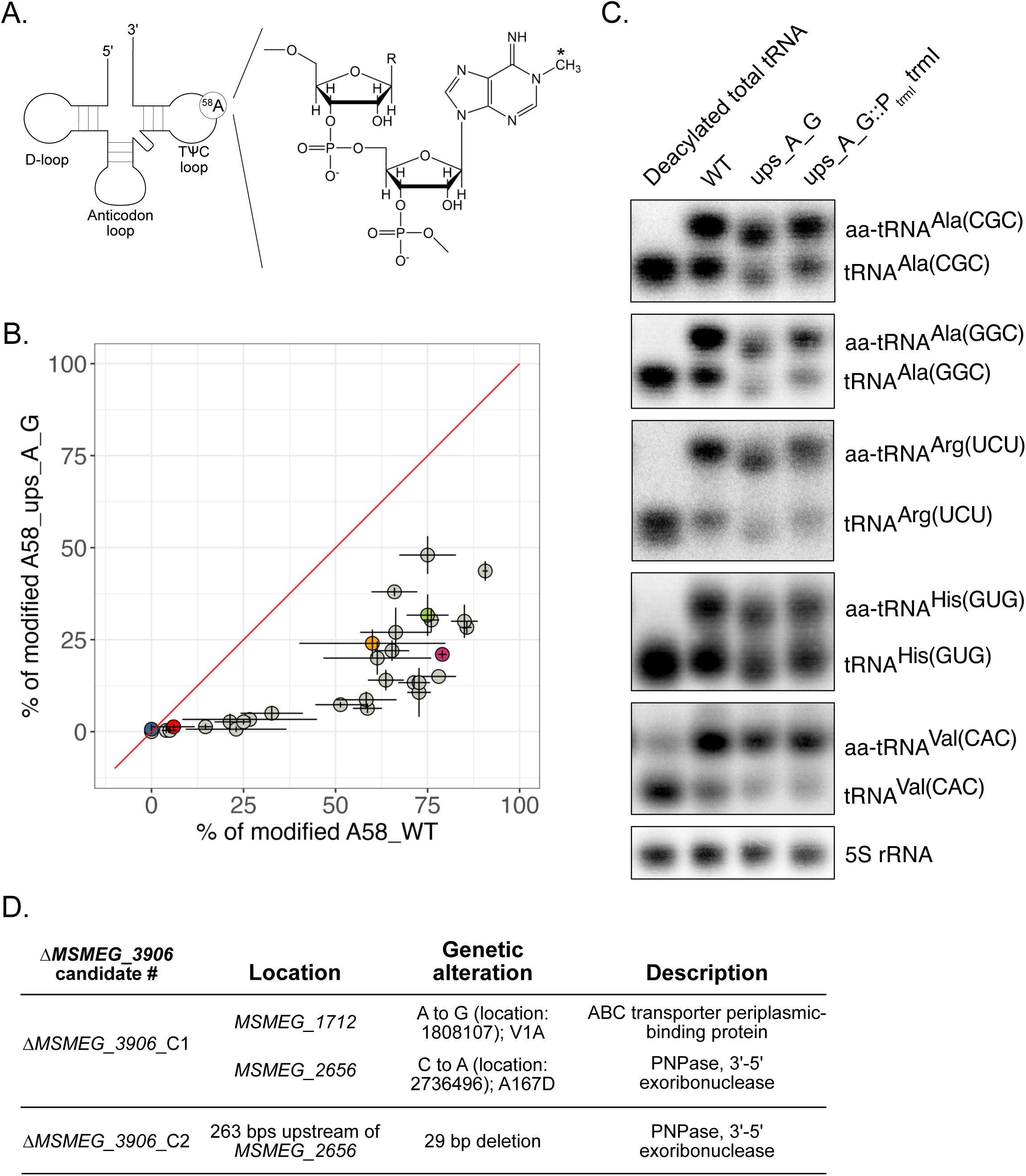
Loss of function of MSMEG_3906 leads to hypomodification of tRNAs, thereby impacting tRNA stability. **A.** Schematic representation of a tRNA. The D-loop; the anticodon loop, which recognizes cognate codons; and the TΨC loop are depicted. The adenine at position 58 (^58^A) is conserved in all *Msm* tRNAs and is methylated (*) by the adenine-N(1)-methyltransferase TrmI (right side of figure). **B.** Quantification of modified adenine at position 58 in WT *Msm* and the ups_A_G adenine-N(1)-methyltransferase mutant using small RNA sequencing. Methylated adenine is mis-read as another base during sequencing, so the level of modification can be determined by comparing the rate of mis-reading at position 58. The red line represents an equal level of A58 modification between the two strains, and each circle represents a tRNA. Error bars represent +/− SD. Circles that fall below the red line indicate tRNAs for which adenine at position 58 is modified more frequently in WT *Msm* than in the mutant. tRNA^Ala(GGC)^, red; tRNA^Arg(UCU)^, green; tRNA^His(GUG)^, pink; tRNA^Ala(CGC)^, orange; tRNA^Val(CAC)^, blue. **C.** Northen blots run on denaturing acid polyacrylamide gels to detect charged (aminoacyl-tRNA, aa-tRNA) and uncharged tRNAs. The degree of methylation affects the distance the tRNA travels in the gel for both charged and uncharged tRNAs. A shift relative to the WT (WT) is observed for Ala, Arg, and His tRNAs from the ups_A_G *trml* mutant but not for tRNA^Val^. Complementing the ups_A_G *ms*_*trml* mutant with a WT copy of *trml* corrects the shifts. **D.** Additional genetic alterations identified in the two *MSMEG_3906* (*trml*) deletion mutants we obtained. Both mutants, for which deletion of *trmI* was confirmed via whole-genome sequencing, had additional mutations related to the gene *MSMEG_2656*, which encodes a 3’-5’ exoribonuclease polynucleotide phosphorylase (PNPase).

The ups_A_G mutant retains about 15% of *ms_trmI* expression compared to the WT (Fig. 1E). To more cleanly investigate the role of ms_TrmI in high survival, we attempted to construct a *MSMEG_3906* deletion mutant. Numerous attempts using multiple methods yielded only two mutants (Fig. 2D; Supplemental Fig. 2B), which was unexpected in light of previous reports that showed that while knocking down *ms_trmI* produced a severe fitness cost (https://pebble.rockefeller.edu/genes/MC2-155/MSMEG3906/), the gene was not essential (https://msrdb.org) (Bosch *et al*., 2021; Judd *et al*., 2021). Unlike the ups_A_G and G128S mutants, the two *ms_trmI* deletion mutants we obtained did not display high survival to kanamycin (Supplemental Fig. 2B). Whole-genome sequencing of the two candidate mutants revealed that they both had mutations related to the essential gene *MSMEG_2656*, which encodes the 3’-5’ exoribonuclease polynucleotide phosphorylase (PNPase), part of the RNA degradosome (Fig. 2D). A 29-bp deletion upstream of the PNPase coding sequence in one of the mutants led to decreased expression levels (Supplemental Fig. 2C). Our inability to recover *ms_trmI* deletion mutants without additional mutations suggested that stabilization of tRNAs by ms_TrmI-mediated methylation is essential for *Msm* viability. Further, the ability of mutations leading to partial loss of PNPase function to rescue *ms_trmI* deletion mutants was likely due to compensatory stabilization of tRNAs in those mutants. These results demonstrate that both *trmI* mutations induce a chronic shortage of functional tRNA through destabilization of the tRNA pool. Next, we aimed to identify the molecular mechanisms by which hypomethylation-driven tRNA instability leads to high survival upon exposure to aminoglycosides.

### tRNA instability leads to increased levels of the transcriptional regulator WhiB7

Using transcriptomics, we found that the ups_A_G mutant displayed increased expression of *whiB7* and members of its regulon, including *eis*, which encodes an acetyltransferase that inactivates aminoglycosides (Fig. 3A, green; Table S1). The transcriptional signature of the ups_A_G isolated mutant overlapped substantially with those of our previously published *argA* and *argD* mutants, in which perturbations in the arginine biosynthesis pathway led to upregulation of *whiB7* (Fig. 3A, circled in red). To test if WhiB7 and Eis mediated increased survival upon exposure to aminoglycosides in the *ms_trmI* high-survival mutants, we deleted either *whiB7* or *eis* in the ups_A_G and G128S mutants. Deletion of either gene led to decreased survival, supporting a model wherein tRNA hypomodification leads to increased expression of the transcriptional regulator WhiB7 and members of its regulon, including Eis, which directly mediates high survival upon exposure to aminoglycosides (Fig. 3B, Supplemental Fig. 3A).

**Figure 3.**
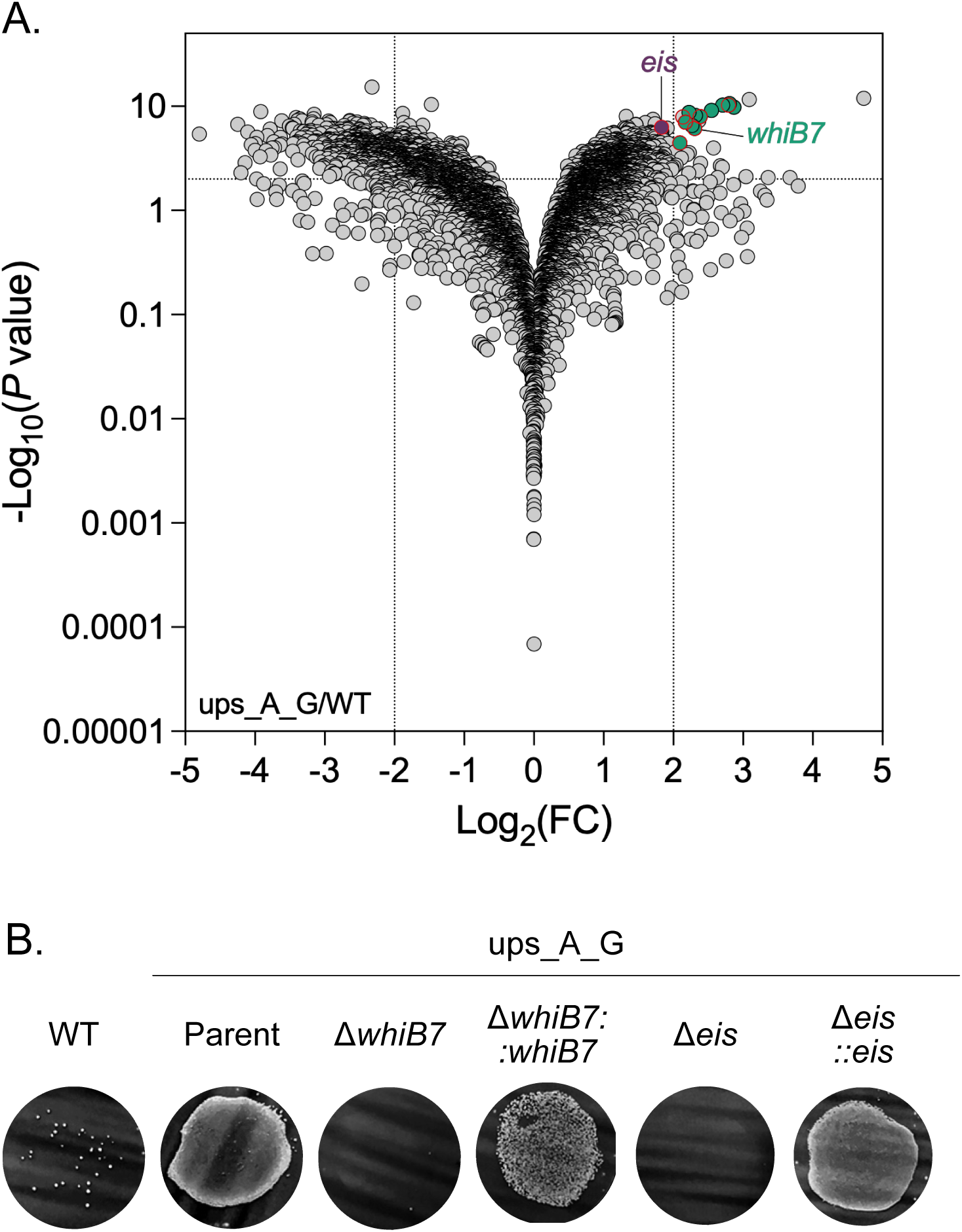
Loss of a tRNA epitranscriptional modification leads to increased levels of the transcriptional regulator WhiB7. RNA sequencing was performed on RNA extracted from ups_A_G and WT (WT) strains of *Msm.* **A.** −Log10 P value versus log2 fold change (FC) compared to WT for each gene (symbols). Dotted lines mark biologic (FC +/− 4) and statistical (P value 0.05) significance cutoffs. Green symbols indicate genes that belong to the WhiB7 operon; those that overlap with genes upregulated in the previously reported *ΔargD* and *ΔargA* mutants (Schrader *et al*., 2021) are outlined in red. Eis (purple) also belongs to the WhiB7 operon and encodes an acetyltransferase that inactivates aminoglycosides. **B.** Kanamycin survival of ups_A_G, ups_A_G_Δ*whiB7*, ups_A_G_Δ*eis*, and the respective complemented strains, in which expression of the complementing gene is under control of its native promoter. Representative of four and three independent experiments for Δ*whiB7* and Δ*eis*, respectively.

The region directly upstream of the *whiB7* gene mediates a transcriptional attenuation mechanism that involves a short upstream open reading frame (uORF) (Burian *et al*., 2012; Dinan *et al*., 2014; Lee, Lee and Roe, 2022). Under non-stress conditions, the rate at which the uORF is translated favours the formation of a Rho-independent terminator that limits *whiB7* transcription. Under stress conditions, including exposure to antibiotics that affect proteostasis, ribosome stalling slows translation of the uORF, allowing it to adopt an antiterminator formation that facilitates RNA polymerase readthrough and transcription of *whiB7* (Lee, Lee and Roe, 2022; Poulton *et al*., 2024). WhiB7 then facilitates transcription of members of its regulon, some of which are known to be involved in antibiotic recalcitrance and resistance (Fig. 4A). Collectively, our results showed that in the ups_A_G mutant, lack of tRNA methylation at position A58 leads to tRNA instability while preserving tRNA charging. A shortage of tRNAs would promote ribosome stalling in the uORF of *whiB7*, thereby inducing expression of *whiB7* and members of its regulon (Fig. 4A). However, upon proteotoxic stress, stalled ribosomes are recycled to the pool of ribosomes available for translation via an essential mechanism called *trans-*translation (Keiler, Waller and Sauer, 1996; Keiler, 2008). This process would be expected to suppress WhiB7 expression, posing a conundrum: how can mycobacterial cells simultaneously maintain both robust WhiB7 activation and trans-translation?

**Figure 4.**
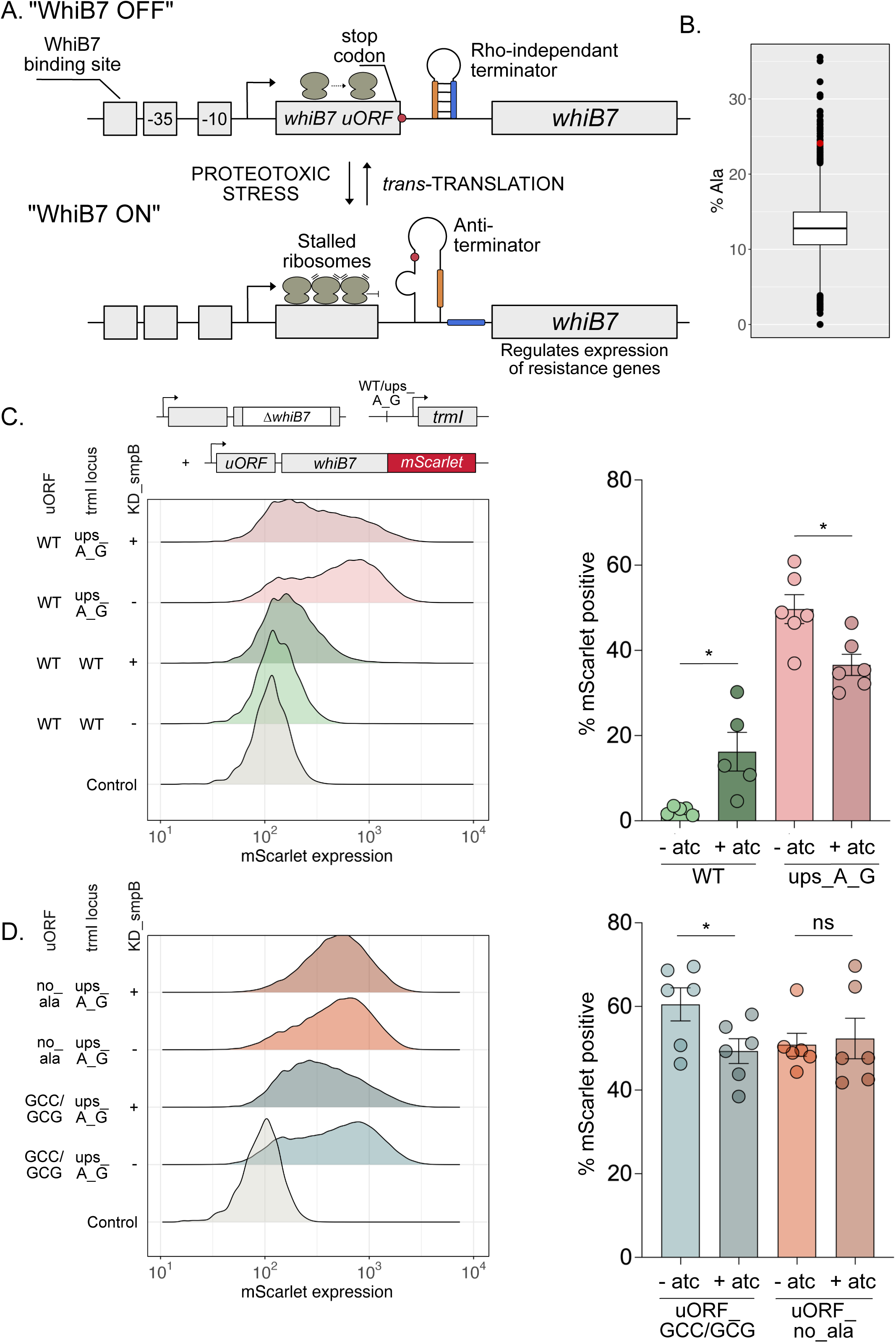
Alanine couples *trans*-translation to WhiB7-mediated riboregulation of resistance genes. **A.** Schematic representation of the *Msm whiB7* locus. The genomic region upstream of *whiB7* contains an upstream open reading frame (uORF) that encodes an 83-amino-acid product. In the absence of stress, the uORF is rapidly translated, promoting formation of a rho-independent terminator that ends transcription and prevents *whiB7* expression (“OFF”). In the presence of stress, especially proteotoxic stress, ribosome stalling slows translation, promoting formation of an anti-terminator structure that permits downstream expression of *whiB7* (“ON”). *trans*-translation is an alanine-dependent process that rescues stalled ribosomes by freeing them from incomplete polypeptides, which it simultaneously targets for degradation. **B.** Box-and-whisker plot (box, interquartile range (IQR) with line at median; whiskers, smallest and largest values within 1.5 × IQR of the first and third quartiles; filled circles, outliers) showing the alanine content of individual *Msm* proteins. The uORF of *whiB7* (red circle) is among the proteins with the highest alanine content. **C.** WhiB7-mScarlet expression measured by flow cytometry in strains of *Msm* (wild-type [WT] and ups_A_G) in which the native allele of *whiB7* has been deleted and *whiB7-mScarlet* has been chromosomally integrated under control of the native *whiB7* promoter with an unmodified uORF. The strains also contain CRISPRi machinery that enables knockdown (KD) of *smpB* upon exposure to anhydrotetracycline (atc). The left panel shows fluorescence intensity peaks at 561 nm/600-620 nm, while the right panel reports the percentage of mScarlet positive bacteria in each condition. WT *Msm* was used as a no fluorescence control. Bars represent mean values of three biologic replicates from six experiments (circles). Error bars represent +/− SD. *, p < 0.05. **D.** Same as (C), except that *whiB7-mScarlet* is expressed under the control of the native *whiB7* promoter with a modified uORF. In one modification, all alanine-coding GCC codons have been replaced with GCG alanine codons (GCC/GCG). In the second, all alanine-coding codons except for the last one, which is essential for the formation of the antiterminator, have been replaced with codons for other amino acids (no_ala). Ns, not significant.

### Alanine levels couple *trans*-translation to WhiB7-mediated riboregulation of antibiotic resistance/recalcitrance genes

In *trans-*translation, a tmRNA—which has properties of both a tRNA and an mRNA—folds into a structure resembling Ala-tRNA to allow recognition by alanyl-tRNA synthetase (AlaRS), which charges its 3’ end with alanine. An accessory protein called SmpB binds to tmRNA with high affinity and acts to stabilize its structure and facilitate its functions. The tmRNA-SmpB complex binds to and rescues stalled ribosomes by transferring them to its mRNA-like domain (MLD); translation of the short MLD results in addition of an alanine-rich tag to the incomplete polypeptides stuck in the stalled ribosomes, targeting the polypeptides for degradation by the ClpC pathway and freeing the ribosomes when the MLD stop codon is reached.

We reasoned that to cope with stress conditions, particularly antibiotic-mediated proteotoxic stress, *Msm* requires both WhiB7-mediated expression of antibiotic resistance and recalcitrance genes and active *trans-* translation. However, it seemed that rescue of stalled ribosomes via *trans*-translation would stymie upregulation of *whiB7* given its mechanistic reliance on ribosome stalling. To reconcile this conundrum, we hypothesized that *trans-*translation might consume a molecule whose shortage would specifically increase ribosome stalling in the uORF of *whiB7* and thus facilitate *whiB7* expression even in the presence of intact *trans*-translation. Notably, the amino acid alanine comprises 24.1% of the uORF of *whiB7*, among the highest alanine content in the *Msm* proteome (Fig. 4B & Table S2). Since *trans-*translation is an alanine-dependent process, we reasoned that consumption of alanine or demand for alanyl-tRNAs in translation of the alanine-rich tag added to prematurely terminated polypeptides during *trans-*translation could selectively increase ribosome stalling in the alanine-rich uORF of *whiB7*. To evaluate if *trans-*translation had an effect on WhiB7 expression, we expressed a functional, mScarlet-tagged WhiB7 protein under the control of its native promoter in *ms_trmI* WT, Δ*whiB7* and ups_A_G, Δ*whiB7* strains equipped with a CRISPR interference (CRISPRi) system capable of partially blocking *trans-*translation via knockdown of the tmRNA co-factor SmpB. We then measured mScarlet expression levels by flow cytometry in the presence and absence of SmpB knockdown (Supplemental Fig. 4A). First, in line with our hypothesis, partially blocking *trans-*translation led to decreased WhiB7 expression in the ups_A_G mutant, in which *whiB7* expression is constitutively increased compared to WT *Msm* (Figure 4C; light pink vs. light green; Supplemental Fig. S4A), but increased WhiB7 expression in WT *Msm* (Fig. 4C). This suggested that in addition to tRNA shortage, *trans-*translation also contributed to increased WhiB7 expression in the ups_A_G mutant.

The alanine-rich tag encoded by the MLD of *Msm* tmRNA uses primarily the GCC codon for alanine. To test if *trans-*translation was depleting the pool of alanyl-tRNA^CGG^, we substituted an alternative alanine codon for all the GCC alanine codons in the uORF of *whiB7* (see methods) and demonstrated that the construct was able to mediate high survival upon exposure to kanamycin when transformed into the ups_A_G mutant in which WT *whiB7* had been deleted, supporting retained functionality (Supplemental Fig. 4B & 4C). We reasoned that if shortage of alanyl-tRNA^CGG^ is sensed by the uORF of *whiB7*, substitution of GCC codons for alternative alanine codons should make it insensitive to alanyl-tRNA^CGG^ shortage, and knocking down *trans-*translation should no longer lead to a decrease in WhiB7 expression. However, we still observed a decrease in WhiB7 expression upon knocking down *trans*-translation (Fig. 4D), suggesting alanyl-tRNA^CGG^ shortage was not sensed by the *whiB7* uORF. We next tested whether alanine consumption during *trans-*translation contributed to increased *whiB7* expression in the ups_A_G mutant. To do so, we substituted other amino acids for all alanine residues in the uORF of whiB7 except one for which the associated codon is critical for the formation of the antiterminator. This modified construct was able to mediate a high survival phenotype in the ups_A_G mutant with WT *whiB7* deleted, demonstrating that it was functional (Supplemental Fig. 4B & 4C). Knocking down *trans-*translation in the ups_A_G mutant containing *whiB7* with the alanine-poor uORF no longer altered WhiB7 expression (Figure 4D). This suggested that *trans-*translation was no longer contributing to WhiB7 expression levels via consumption of alanine and increased ribosome stalling on the *whiB7* uORF. Collectively, these results showed that alanine couples *trans-*translation to riboregulation of WhiB7-mediated resistance and recalcitrance genes. These findings reveal that the high survival phenotype of *ms_trmI* mutants depends on sophisticated metabolic coordination between seemingly competing cellular processes.

## DISCUSSION

Tolerance and persistence allow bacteria to survive otherwise lethal concentrations of an antibiotic. They cause prolonged treatment, contribute to infection relapse, and are harbingers of antibiotic resistance (Levin-Reisman *et al*., 2017; Sebastian *et al*., 2017; Etthel M. Windels *et al*., 2019; Huemer *et al*., 2020; Liu *et al*., 2020), providing an incentive for development of therapeutic strategies that kill tolerant and persistent bacteria. However, this requires a deeper understanding of the molecular mechanisms underlying tolerance and persistence. Such knowledge would also inform the development of diagnostic tools to evaluate antibiotic recalcitrance in clinical settings.

Towards this goal, we developed the ASSIST method for selection and isolation of antibiotic high-survival mutants and previously used it to isolate 29 mutants in *Msm* (Schrader *et al*., 2021). The two mutants studied here displayed high survival upon exposure to aminoglycosides and macrolides due to alterations affecting a gene encoding a tRNA methyltransferase. tRNA modification has been implicated in bacterial response to antibiotic stress (de Crécy-Lagard and Jaroch, 2021). For example, the toxin TactT acetylates the primary amine group of amino acids on charged tRNA molecules in *Salmonella*, thereby inhibiting translation and promoting persister cell formation (Cheverton *et al*., 2016). In addition, loss of function of tRNA-modifying enzymes was shown to alter antibiotic susceptibility in *Vibrio cholerae* (Babosan *et al*., 2022; Fruchard *et al*., 2025), and bacteria lacking m¹G37 were shown to have impaired membrane structure, rendering them more susceptible to multiple classes of antibiotics (Masuda *et al*., 2019; Hou, Masuda and Foster, 2020). Here, we found that hypomethylation of multiple tRNAs in *ms_trmI* mutants led to tRNA instability and upregulation of the transcriptional regulator WhiB7. In contrast to the activity of Mtb_TrmI, which was restricted to formyl-methionine tRNA when heterologously expressed in *E. coli* (Varshney *et al*., 2004), we found that the *Msm* tRNA-modifying enzyme adenine-N(1)-methyltransferase ms_TrmI could modify a broad range of tRNAs. We also found that we could not knock out *ms_trmI* unless concomitant mutations were present in the gene encoding PNPase. Our inability to isolate *ms_trmI* knockouts without concomitant PNPase mutations suggests that *ms_trmI* is essential in *Msm*, likely because tRNAs lacking *ms_trmI*-mediated methylation are unstable and rapidly degraded. Given that PNPase is essential in *Msm*, the mutations we found likely resulted in partial loss of PNPase function, which reduced degradation of tRNAs and prevented cell death. This aligns with prior studies in *S. cerevisiae*, where spontaneous suppressor mutations arose in the 3′-5′ exoribonuclease *DIS3/RRP44* to compensate for initiator tRNA hyperinstability caused by mutations in *trm6*, a subunit of *trmI* (Anderson et al., 1998; Kadaba et al., 2004). Notably, while we were able to delete the ms_trmI homolog rv2118c in *Mycobacterium tuberculosis* (*Mtb*), this did not lead to high survival upon exposure to aminoglycosides. This result could reflect restriction of mtb_TrmI activity to formyl-methionine tRNA in concordance with the results of heterologous expression experiments in *E. coli* (Varshney *et al*., 2004) or could alternatively suggest that other redundant methyltransferases or compensatory mechanisms exist in *Mtb* (Supplemental Fig. 5).

In addition to examining the function of *ms_trmI* in *Msm*, we explored how mutations leading to reduced ms_*TrmI* expression conferred increased survival upon exposure to aminoglycosides. Our data indicate that reduced tRNA methylation and subsequent global tRNA shortage activate the transcriptional regulator WhiB7 because of increased stalling on the uORF of *whiB7*, which likely serves as a sensor of proteotoxic stress in mycobacteria (Poulton *et al*., 2024). Activation of WhiB7, in turn, induces the expression of Eis, an acetyltransferase that inactivates aminoglycosides. Further, we link *trans-*translation to regulation of WhiB7 expression and recalcitrance mediators. *Trans-*translation is an essential process that allows bacteria to cope with ribosome stalling, a phenomenon that occurs at low levels in non-stress conditions and amplifies upon proteotoxic stress. Although activation of *trans*-translation counteracts ribosome stalling on a global level, our results demonstrate that alanine consumption during synthesis of alanine-rich tags used to mark incomplete peptides during ribosome rescue specifically intensifies ribosome stalling on the alanine-rich uORF of *whiB7.* Since *whiB7* expression is dependent on ribosome stalling, which promotes formation of the antiterminator structure in the uORF, this selective intensification serves to protect *whiB7* expression from the ribosome rescuing effects of *trans-*translation that would otherwise decrease it. Our data also show that intensification of ribosome stalling by alanine depletion not only maintains but also contributes to upregulation of *whiB7* and the recalcitrance mediating genes it regulates, as an uORF construct with most alanine residues removed produced a lower level of WhiB7 expression than a construct retaining these residues. Notably, alanine aminotransferase—a key enzyme in the alanine biosynthesis pathway in *Msm*—belongs to the *whiB7* operon. We speculate that WhiB7-mediated upregulation of alanine aminotransferase facilitates replenishment of alanine levels in a positive feedback loop that sustains both *trans-*translation and WhiB7 expression to combat proteotoxic stress (Fig. 5). Given the key role of the uORF structure in mediating *whiB7* expression, this represents an example of riboregulation of bacterial tolerance genes, extending the findings of others who have described the control of antibiotic resistance genes by riboregulators (Vazquez-Laslop, Thum and Mankin, 2008; Dar *et al*., 2016; Dersch *et al*., 2017).

**Figure 5.**
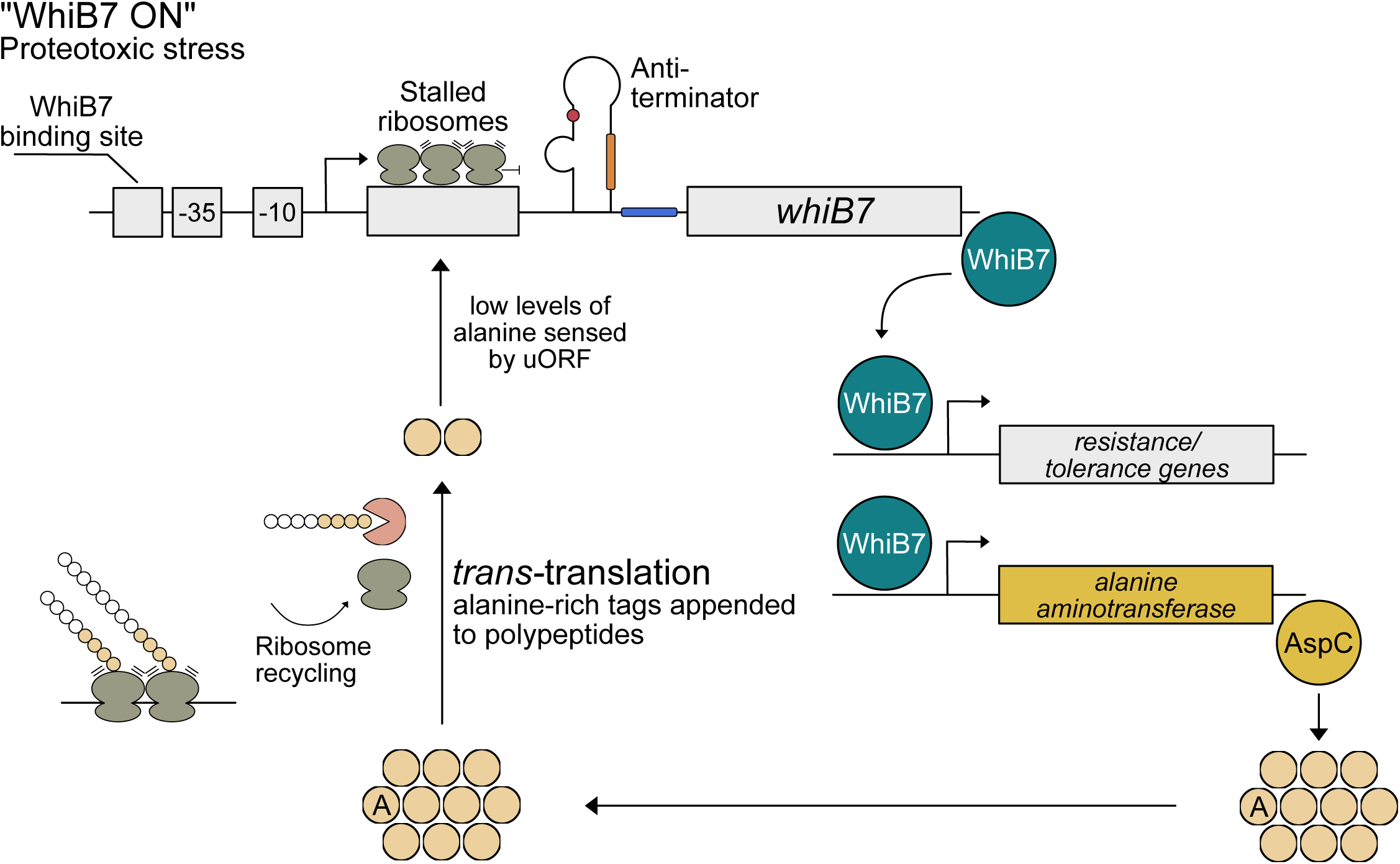
Schematic representation of the mechanism by which alanine couples *trans-*translation to riboregulation of WhiB7-mediated tolerance genes. Upon proteotoxic stress, ribosome stalling in the uORF of *whiB7* promotes the formation of an antiterminator so that *whiB7* is expressed, leading to upregulation of members of its regulon. At the same time, *trans*-translation is activated in response to increased ribosome stalling. Alanine consumption by *trans*-translation specifically amplifies ribosome stalling on the alanine-rich uORF of *whiB7*. This mechanism allows for simultaneous expression of WhiB7 and *trans*-translation to cope with proteotoxic stress. The WhiB7 regulon includes the alanine aminotransferase AspC, which likely permits replenishment of the alanine pool to sustain *trans*-translation.

We previously characterized the role of WhiB7 in mediating increased survival upon exposure to aminoglycosides in *Msm* strains with mutations in the arginine biosynthesis pathway. Although the mechanisms by which defects in the arginine biosynthesis pathway led to increased expression of *whiB7* are unknown, given our findings here, it is tempting to speculate that a polyarginine stretch in the uORF of whiB7 (Table S2) might promote ribosome stalling and antiterminator formation in those mutants. In addition to perturbation of arginine biosynthesis, the demonstration here of another pathway—tRNA methylation—that leads to upregulation of the transcriptional regulator WhiB7 and subsequent resistance and recalcitrance upon exposure to antibiotics supports the classification of WhiB7 as a molecular convergence point in the control of multiple mechanisms of antibiotic resistance and recalcitrance. Work by others also lends support to this conclusion (Morris *et al*., 2005; Burian *et al*., 2012, 2013; Hurst-Hess, Rudra and Ghosh, 2017; Schildkraut *et al*., 2022; Troian *et al*., 2023; Bernard *et al*., 2024). For example, a *Mycobacterium abscessus* (*Mab*) mutant with a defect in the serine biosynthesis pathway overexpressed WhiB7 and was tolerant to cefoxitin, a β-lactam antibiotic (Bernard *et al*., 2024). In another example, cleavage of tRNA^Ser(CGA)^ by the *Mab* VapC5 toxin led to WhiB7 overexpression and tolerance to amikacin, tedizolid, and cefoxitin (Troian *et al*., 2023). In both cases, it is conceivable that serine or tRNA^Ser^ shortage led to ribosome stalling, thus triggering *trans*-translation. In accordance with our findings, alanine consumption by *trans*-translation might then activate WhiB7 and the antibiotic resistance/recalcitrance genes in its regulon by increasing ribosome stalling in the alanine-rich *whiB7* uORF, which is well-conserved among mycobacterial species, including *Mab* and *Mtb*. Collectively, our findings suggest that mycobacterial species might have evolved a protection mechanism, perhaps representing a unique adaptation in *Actinomycetales*, that supports both *trans-*translation and expression of tolerance and resistance genes upon exposure to proteotoxic stress.

Tolerance and persistence arise through numerous pathways; developing a drug to target each individual mechanism is both financially and therapeutically impractical. Therefore, identifying key regulatory nexuses—such as WhiB7—that link multiple tolerance and persistence mechanisms is a crucial step toward developing effective intervention strategies.

---

**Supplemental figure 1.**
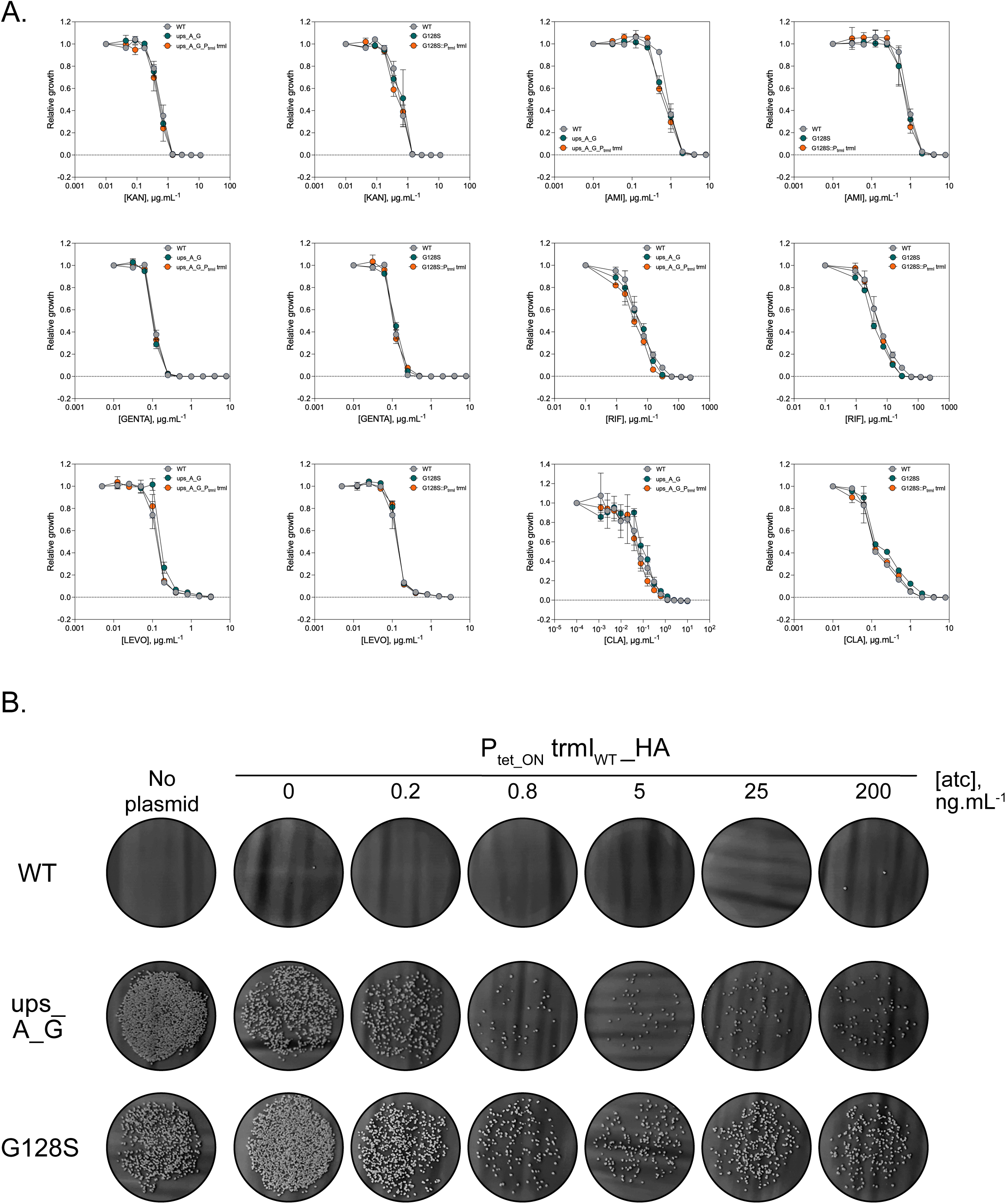
**A.** Kanamycin (KAN), amikacin (AMI), gentamycin (GENTA), rifampicin (RIF), levofloxacin (LEVO) and clarithromycin (CLA) MICs for WT (WT) *Msm*, the ups_A_G and G128S mutants, and their respective complemented strains. Growth is shown relative to a no-antibiotic control. Symbols, means of three replicates. Error bars represent +/− SD. Representative of two independent experiments. **B.** Survival of WT *Msm*, the ups_A_G and G128S mutants, and their respective complemented strains after exposure to kanamycin. Mutants were complemented with an anhydrotetracycline-inducible allele (Ptet_ON) encoding WT ms_TrmI with an HA tag at the C-terminus; the ability of TrmI-HA to effectively complement the high-survival phenotype of the ups_A_G and G128S mutants demonstrates that the construct is functional.

**Supplemental figure 2.**
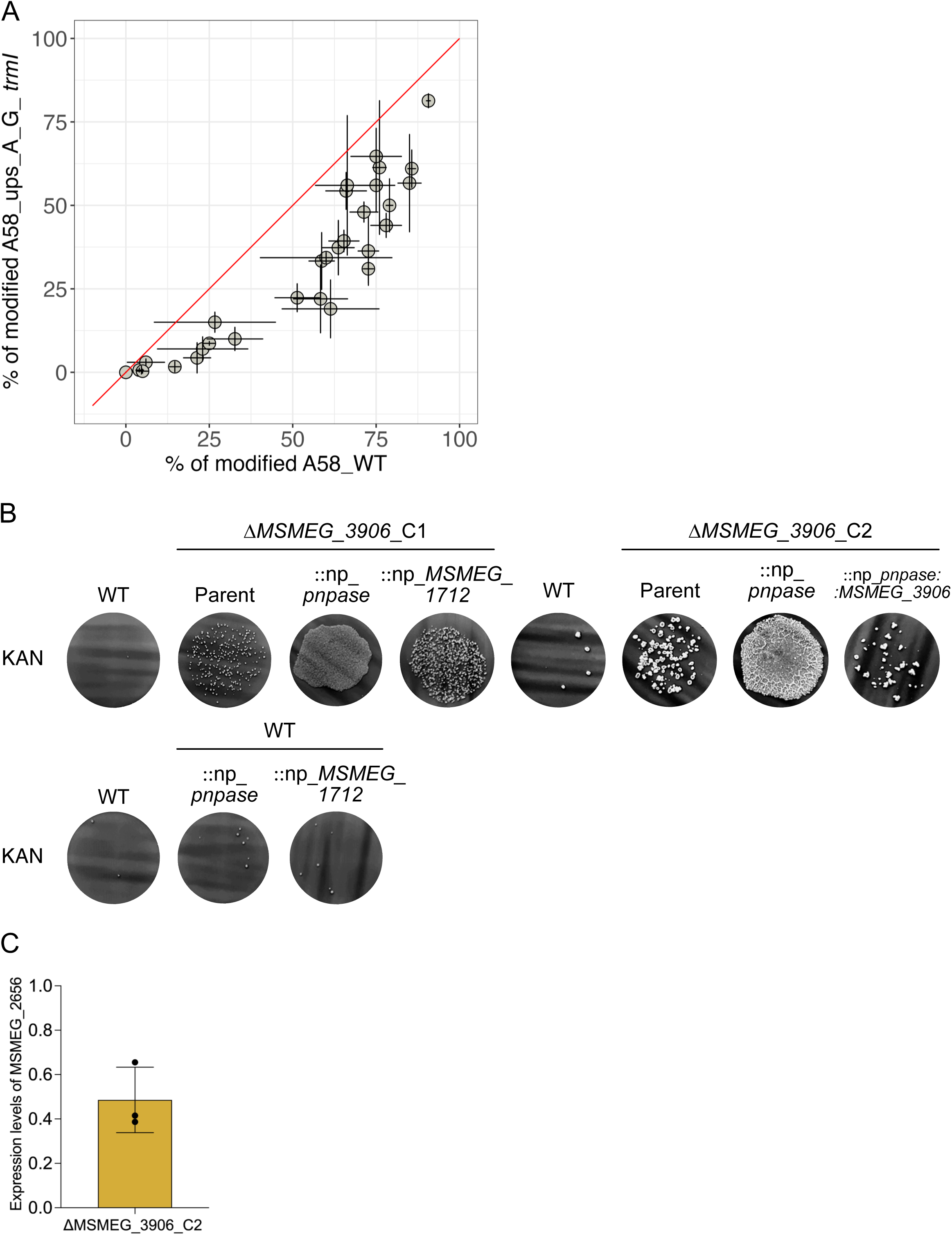
**A.** Quantification of modified A^58^ in the ups_A_G adenine-N(1)-methyltransferase mutant complemented with a WT trml allele using small RNA sequencing. The red line represents an equal level of A58 modification between the two strains, and each circle represents a tRNA. Error bars represent +/− SD. **B.** Complementation of the *ΔMSMEG_3906* deletion mutants (C1 and C2) with WT versions of other genes in which mutations were found after mutant construction, each under control of its native promoter (np). Addition of the gene encoding PNPase or the gene encoding MSMEG_1712 in WT *Msm* does not confer high survival upon exposure to kanamycin (KAN). **C.** Expression level of *MSMEG_2656* (encoding PNPase) in *ΔMSMEG_3906* candidate #2 (C2) compared to WT *Msm*. The bar represents mean expression level across three biological replicates (black circles). Error bars represent +/− SD.

**Supplemental figure 3.**
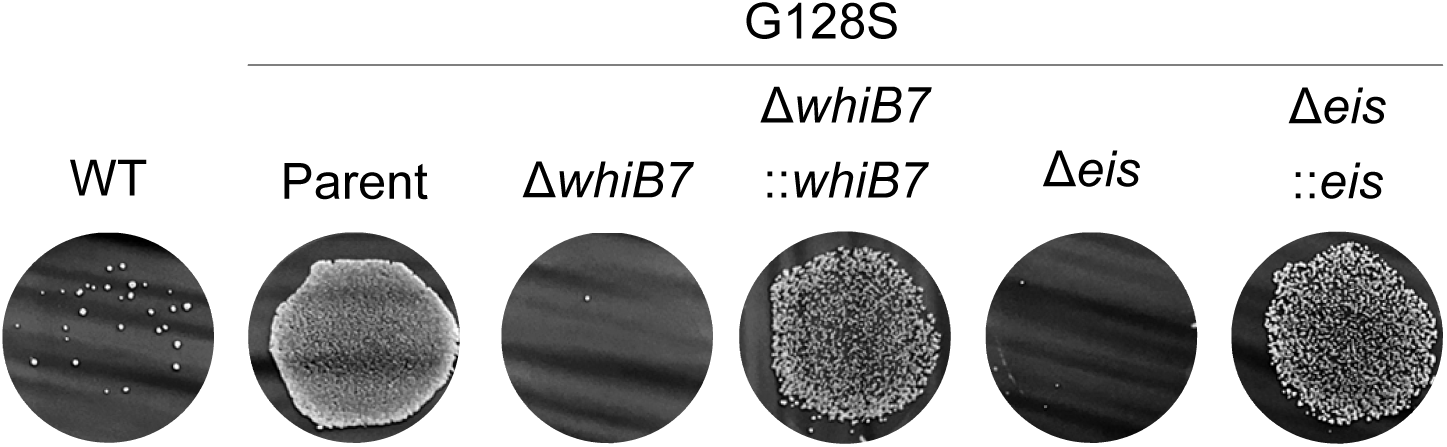
Kanamycin survival of G128S, G128S_Δ*whiB7*, G128S_Δ*eis*, and G128S_Δ*whiB7*, G128S_Δ*eis* complemented with either *whiB7* or *eis*. Representative of four independent experiments.

**Supplemental figure 4.**
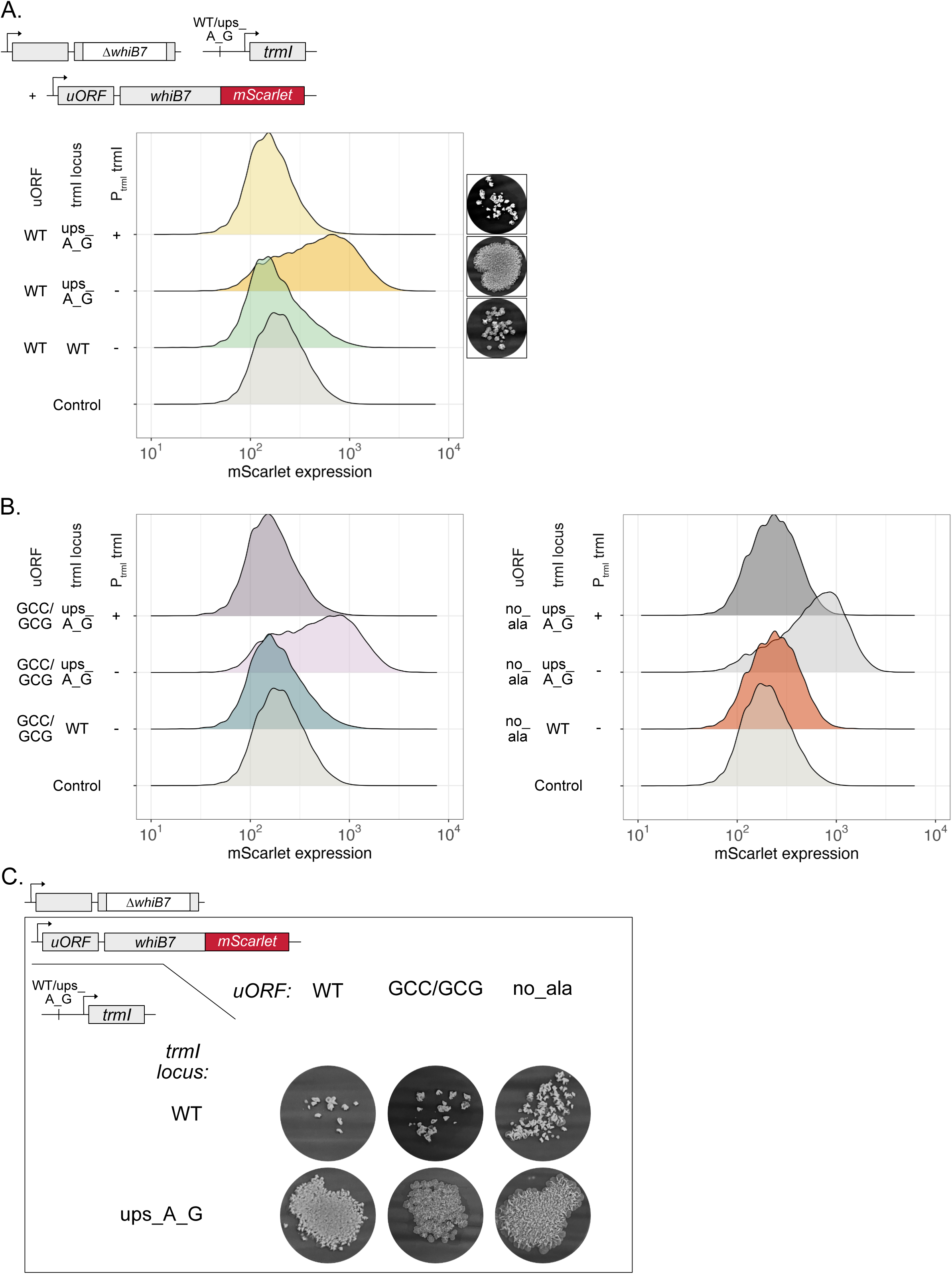
**A.** WhiB7-mScarlet expression in strains of *Msm* (Wild type [WT] and the ups_A_G) in which the native allele of *whiB7* has been deleted and *whiB7-mScarlet* is chromosomally integrated under the control of the native whiB7 promoter with an unmodified uORF. The A-to-G mutation in the promoter of *ms_trmI* leads to increased expression of WhiB7-mScarlet and increased survival upon exposure to kanamycin (photographs of filters shown to the right of the corresponding flow plots); WhiB7-mScarlet expression and high survival upon exposure to kanamycin are complemented by expression of *ms_trml* under its native promoter. **B and C**. Same as (A), except that *whiB7-mScarlet* is expressed under the control of the native *whiB7* promoter with a modified uORF. In one modification, all alanine-coding GCC codons have been replaced with GCG alanine codons (GCC/GCG). In the second, all alanine-coding codons except for the last one, which is essential for the formation of the antiterminator, have been replaced with codons for other amino acids (no_ala) shows mScarlet expression measured by flow cytometry, while C) shows survival upon exposure to kanamycin. The results shown demonstrate that constructs containing a modified uORF are functional because like the unmodified uORF, they allow for high survival upon exposure to kanamycin.

**Supplemental figure 5.**
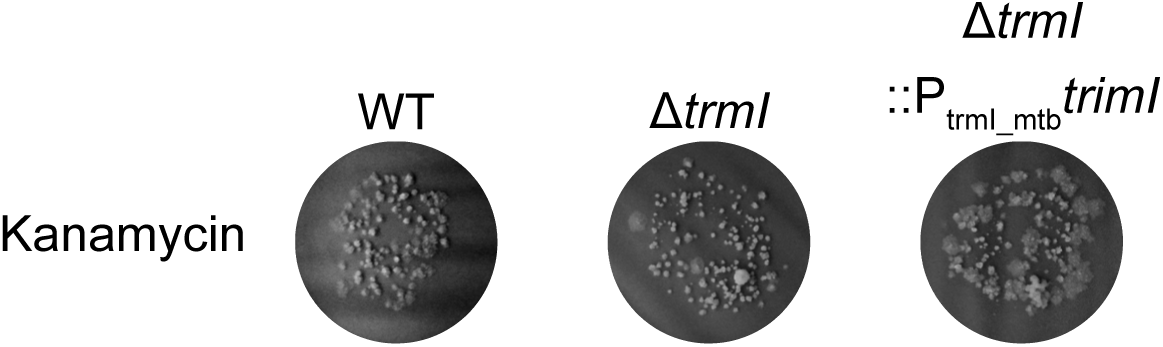
Survival upon exposure to kanamycin of WT Mtb, Δ*trmI*, and Δ*trmI*::P_­­­­_::*trmI*. Representative of two independent experiments.

## MATERIALS AND METHODS

### Bacterial strains and culture conditions

*Msm* strains were cultured at 37°C with shaking in Middlebrook 7H9 medium (BD Difco) supplemented with 0.5% bovine serum albumin Fraction V (Roche), 0.2% glycerol, 0.2% dextrose, 0.05% tyloxapol, and 0.085% sodium chloride. Appropriate antibiotics [hygromycin (50 µg.mL^−1^, Roche), zeocin (25 µg.mL^−1^, Thermo Fisher Scientific), streptomycin (25 µg.mL^−1^, Sigma-Aldrich), or kanamycin (25 µg.mL^−1^, Thermo Fisher Scientific)] were added during generation of strains bearing an episomal or chromosomally integrated antibiotic resistance cassette and excluded in cultures grown for experiments with the successfully generated strains. For experiments requiring knockdown of SmpB, *Msm* strains containing the pLJR962 plasmid for CRISPR interference (CRISPRi) were cultured in 7H9 medium supplemented with 20 ng.µL^−1^ of atc for 16 hours prior to flow cytometry analysis. All strains used in this study are listed in Table S3.

### Plasmid construction

Except for the plasmids bearing sgRNAs used for CRISPRi, all plasmids were constructed using Invitrogen Gateway recombination cloning technology. The plasmids were transformed into competent *E. coli* DH5α cells or *E. coli* MACH1 cells (Thermo Fisher Scientific). Primers used in cloning are listed in Table S3. For CRISPRi, the *Msm*-adapted, atc-regulated CRISPRi backbone (pLJR962) with a catalytically inactivated CRISPRi cas9 allele from *Streptococcus thermophilus* was kindly provided by Dr Jeremy Rock (The Rockefeller University). The sgRNA targeting Msm *smpB* (*MSMEG_2091*) was designed following the guidelines described in Rock et al. (Rock *et al*., 2017), with the protospacer adjacent motif (PAM) determined based on the published consensus to obtain the strongest knockdown. To design the sgRNA, 20 nucleotides were added to the 5’ end of the PAM sequence, and 5’-GGGA-3’ and 5’-AAAC-3’ were added to the forward and reverse primers, respectively. The primers were annealed in an annealing buffer at 95°C for 2 minutes followed by a gradual decrease in temperature of 0.1°C/sec to 25°C and then cloned into the pLJR962 backbone after digestion by the restriction enzyme BsmBI (NEB) via ligation by a T4 DNA ligase (NEB). The final plasmid was confirmed by sequencing using the primer 1834 (5’-TTCCTGTGAAGAGCCATTGATAATG-3’).

### whiB7 expression reporter construction

The coding sequence of the fluorescent reporter protein mScarlet was fused to the open reading frame of *Msm whiB7*. To do so, the stop codon of *whiB7* was removed and a linker was added between the *whiB7* and *mScarlet* coding sequences (5’-TCGGGCTCGGGCTCGGGC-3’, encoding the SGSGSG amino acid sequence). We constructed several versions of the *whiB7* promoter (encompassing the 500 bp region upstream of *Msm whiB7*, which includes the uORF to control *whiB7-mScarlet* expression. In the WT construct, we did not modify the uORF from its native sequence. In the GCC-to-GCG construct, all GCC codons in the uORF were replaced with a GCG codon; both codons encode an alanine residue. In the no_ala construct, all alanine-coding codons were replaced with codons for other amino acids except for the last GCG codon, which was retained so as not to perturb the structure of the antitermination loop (see table S2). Altered uORF sequences were synthesized by Genewiz (Azenta Life Sciences).

### Filter seeding for assessment of survival upon exposure to an antibiotic

Liquid cultures grown to exponential phase were diluted to a final concentration of 10^7^ bacteria per mL. One mL of diluted culture was deposited onto a Durapore membrane filter (Merck Millipore, 0.22 µm pore) mounted on a sterilised Swinnex 47 mm filter holder (Merck Millipore) attached to a vacuum canister (Cardinal Health). Seeded filters were transferred onto 7H10 (Middlebrook) plates containing an antibiotic at 10x the MIC and incubated at 37°C for an exposure time pre-determined to kill >99% of a WT bacterial population. Filters were then transferred onto 7H10 plates containing 0.4% charcoal and further incubated for 24h at 37°C to absorb residual antibiotic from the filters. Finally, the filters were placed onto growth-permitting 7H10 plates and incubated at 37°C until colonies appeared (three to four days). Photographs of filters were taken after recovery from antibiotic exposure. Filter images are shown in grayscale with adjusted brightness and contrast.

### Determination of minimum inhibitory concentrations (MICs)

MICs were determined by the standard microdilution method. Briefly, bacterial cultures were subcultured in 7H9 medium and grown to exponential phase at 37°C. Twofold serial dilution series of antibiotics were prepared in the appropriate solvent, and 2 µL of each dilution was added to separate wells of 96-well plates. Bacterial cultures were diluted to an OD_­­­­_of 0.02, and 198 µL of diluted culture per well was added to antibiotic-containing wells. Each dilution was tested in triplicate for each strain. The range of final concentrations tested were: kanamycin, 4.35-1,120 µg.mL^−1^; amikacin, 3.13-800 µg.mL^−1^; levofloxacin, 1.25-320 µg.mL^−1^; rifampicin, 93.75-24,000 µg.mL^−1^; and clarithromycin, 3.13-800 µg.mL^−1^_­­­­_Plates were incubated at 37°C with shaking at 90 rpm for 48 to 72 hours. Wells were mixed to resuspend bacteria before reading the absorbance at 600 nm. The MIC of each antibiotic was defined as the concentration that inhibited >95% of bacterial growth compared to a control without antibiotic.

### Quantitative Reverse Transcription PCR (qRT-PCR)

RNA was extracted as described in the RNA extraction section. 500 ng of RNA was used for reverse transcription with the MuLV reverse transcriptase (Invitrogen) in the presence of an RNase inhibitor (Invitrogen) and random hexamers (Invitrogen). Ambion water was substituted for reverse transcriptase in a no-reverse-transcriptase control used to assess for the presence of residual genomic DNA. cDNA was mixed with PowerTrack SYBR Green Master Mix (ThermoFisher) and cycled with quantification on StepOnePlusTM (ThermoFisher). Crossing points (Cp) for each sample were computed using a second derivative analysis (StepOne software, v2.2.2) for the target and control genes. ΔΔCp values were calculated for each strain by subtracting the average ΔCp (which is the the target gene Cp minus the control gene Cp) of at least three biological replicates of the WT strain from the average ΔCp of tested strains. Data were presented as log2 fold-change relative to the control *sigA* (*MSMEG_2758*), calculated as 2-ΔΔCp. All the primers used for qRT-PCR were ordered from Invitrogen and are listed in Table S3.

### RNA extraction for qRT-PCR and RNA/small RNA sequencing

Cultures were grown to mid-log phase in 10 mL of 7H9 medium. Bacteria were washed with the same volume of 4M GTC buffer (containing 50% guanidium thiocyanate, 0.5% N-lauryl sarcosine, and 0.1M of freshly added 2-mercaptoethanol). Bacterial pellets were subjected to bead-beating to break open the cells, and total RNA was extracted using the Trizol-chloroform method (Zymo Research). Extracted RNA was then purified with the Direct-zol RNA miniprep kit (Zymo Research). After elution, RNA was treated with Turbo DNase (Thermo Fisher Scientific) to eliminate residual DNA, and samples were purified again using the Direct-zol RNA miniprep kit (for qRT-PCR samples) or the RNA Clean and Concentrator 25-kit (Zymo Research, for RNA-sequencing samples).

### RNA sequencing

Total RNA was purified as described in the RNA extraction section. Ribosomal RNA (rRNA) was removed using the RiboMinus Transcriptome Isolation Kit for bacteria (Thermo Fisher Scientific), and samples were concentrated with the RNA Clean and Concentrator-25 kit (Zymo Research). RNA-seq libraries were prepared from rRNA-depleted samples using the NEBNext Ultra II Directional RNA Library Prep Kit for Illumina (New England Biolabs) with Illumina Index Primers Sets 1 and 2 (NEB). Libraries were purified with NEBNext Sample Purification Beads from the Library Prep Kit (NEB). Sample quality was checked using a Bioanalyzer (Agilent), and libraries were pooled for Illumina sequencing on an Illumina HiSeq4000 with 75-bp paired-end sequencing. For data processing, reads were trimmed using the Trim Galore software version 0.6.5 (options -paired -illumina) and aligned to the *Msm* genome obtained from Mycobrowser using Kallisto (version 0.46.1) with the --rf-stranded option (Bray *et al*., 2016). Differential expression analysis was done using DESeq2 (version 1.28.0) as previously described (Stapels *et al*., 2018) to compute the log2 fold-change (log2 FC) and adjusted p-values between compared strains. Log2 FC and adjusted p-values were plotted in Prism (version 10).

### Small RNA sequencing

Total RNA was extracted as described in the RNA extraction section. To isolate small RNA, 2 µg of total RNA was processed using the VAHTS Small RNA Library Prep Kit for Illumina with Universal Primer and Index primers (Vazyme). Library selection was performed using VAHTS DNA Clean Beads (Vazyme). Libraries were quantified by qPCR to adjust concentrations for loading onto a sequencing flowcell. Sequencing was performed using a NextSeq550 with a Midoutput flowcell.

### Knockout construction

The genetic knock-out of *MSMEG_3906* was generated by allelic exchange. WT *Msm* was first transformed with an episomal plasmid containing the phage-derived RecET recombinase (pNIT_RecET_SacB_KanR). RecET expression was induced by adding isovaleronitrile at a final concentration of 12 µM (Sigma Aldrich) to a culture grown to mid-log phase. After a three-hour induction period, competent cells were prepared and transformed with 1 µg of a PCR product containing a genomic region coding for an hygromycin resistance cassette flanked by sequences homologous to the ∼500-bp genomic regions immediately upstream and downstream of *MSMEG_3906*. Colonies that grew on 7H10 plates containing 50 µg.mL^−1^ of hygromycin were screened by PCR to check for insertion of the hygromycin resistance cassette in the *MSMEG_3906* locus. Colonies that met this criterium were outgrown in the absence of kanamycin and plated on 7H10 plates containing 10% sucrose to select for bacteria in which the pNIT_RecET_SacB_KanR plasmid was lost. Sucrose-resistant colonies were then screened for their ability to grow on 7H10 plates without but not with 25 µg.mL^−1^ of kanamycin to confirm plasmid loss. Deletion of *MSMEG_3906* was confirmed by whole-genome sequencing, which also allowed for identification of additional mutations acquired by the candidate strains.

### Whole-genome sequencing and SNP identification

Strains were grown to mid-log to stationary phase in 20 mL 7H9, and genomic DNA was isolated via bead-beating, phenol-chloroform extraction, and ethanol precipitation. Genomic DNA was sequenced at MicrobesNG (University of Birmingham). Deletion of *MSMEG_3906* was confirmed by visually examining the reads around the *MSMEG_3906* locus in Integrated Genomics Viewer (Broad Institute). Single nucleotide polymorphisms (SNPs) were identified by aligning the sequences with the NCBI reference sequence NP_008596.1 using a variant calling pipeline for bacterial genomes developed by the Applied Bioinformatics Core at Weill Cornell Medicine, New York (Schrader *et al*., 2021). The pipeline involves adapter trimming using cutadapt (Martin, 2011), read alignment using a BWA package (*Aligning sequence reads, clone sequences and assembly contigs with BWA-MEM – ScienceOpen*, no date), duplicate marking using Picard’s MarkDuplicates (*Picard Tools - By Broad Institute*, no date), haplotype calling using GATK HaplotypeCaller (Van der Auwera *et al*., 2013), and variant annotation using SnpEff (Cingolani *et al*., 2012).

### Western blotting

To isolate protein, 5 mL of mid-log phase cultures were washed with PBS containing 0.02% tyloxapol. Bacterial pellets were resuspended in 500 µL of PBS supplemented with a protease inhibitor cocktail (Roche) and bead-beat three times for 30 seconds with one minute cooling intervals. After centrifugation, the supernatant was transferred to a new tube, and protein concentration was measured using the DC Protein Assay Kit (Biorad). 8 µg of protein from each sample was loaded on a 12% SDS-PAGE gel and separated by electrophoresis. Gels were transferred using the Trans-Blot Turbo Transfect System (Bio-Rad) onto Immobilon-P polyvinylidene difluoride membranes (Millipore). Blocking of the membranes was done for one hour in 5% (w/v) BSA in TBS-T buffer (150 mM NaCl, 100 mM Tris pH 7.4, 1% Tween-20). Membranes were then incubated overnight at 4°C with rotation in a blocking buffer containing a mouse anti-HA primary antibody (Bio Legend 901503, 1:2000). Membranes were washed three times for ten minutes in TBS-T buffer. Next, membranes were incubated in 5 mL blocking buffer containing a goat anti-mouse HRP-linked secondary antibody (Agilent, P0447, 1:2500) for one hour at room temperature. Membranes were washed three times in TBS-T buffer and developed using the ECL plus chemiluminescence reagents (Pierce, ThermoScientific). Membranes were imaged using a Bio-Rad ChemiDoc imaging system.

### Total RNA extraction under acidic conditions

After inoculating 7H9 medium with an overnight culture of *Msm*, cells were grown to an OD_­­­­_of 0.3 at 37°C and harvested by centrifugation at 10,000 g at 4°C for 10 min. Isolation of total RNA was performed under acidic conditions to maintain aminoacylation (Varshney, Lee and RajBhandary, 1991). Briefly, cell pellets were resuspended in 0.5 ml 0.3 M NaOAc at pH 4.5 and 10 mM EDTA, mixed with 0.5 ml glass beads (0.1 mm, BioSpec Products) and acid-saturated phenol:chloroform (24:1; pH 4.3, Millipore Sigma). The mixtures were vortexed for 1 min and allowed to rest on ice for 1 min. This was repeated 14 times. After centrifugation at 21,000 g for 5 min, the aqueous layers were transferred to new tubes and extracted with acid-saturated phenol:chloroform by vortexing for 1 min. Aqueous layers were transferred to new tubes. After precipitation with 2.5 volumes of ethanol at −80°C, RNA was recovered by centrifugation at 21,000 g for 15 min, washed with 70% ethanol containing 10 mM NaOAc at pH 4.5, dried, and resuspended in acid RNA buffer (10 mM NaOAc at pH 4.5, 1 mM EDTA). For the uncharged tRNA control, the RNA was deacylated in 100 mM Tris-HCl (pH 9), 1 mM EDTA at 37°C for 1 h, precipitated with ethanol and resuspended in H_­­­­_O. **Gel electrophoresis**. To assay aminoacylation, 2 µg of total RNA or the uncharged RNA control were mixed with 2x acid RNA loading buffer (9 M urea, 100 mM NaOAc at pH 5, 0.05% bromophenol blue, and 0.05% xylene cyanol) and fractionated in 6.5% polyacrylamide (19:1 acrylamide:bisacrylamide) gels containing 8 M urea and 100 mM NaOAc at pH 4.5 (Varshney, Lee and RajBhandary, 1991). Following electrophoresis at 350 V for 18 h or until the bromophenol blue band migrated 30 cm, the gel between the xylene cyanol and bromophenol blue dyes was excised, soaked in 1x TBE (50 mM Tris, 45 mM boric acid, 1.25 mM EDTA) for 5 min, and transferred to Hybond N+ Amersham Hybond N+ (Cytiva) in 0.5x TBE at 400 mA for 1 h. For fully denaturing gels, 2 µg total RNA was mixed with 2x RNA loading buffer (95 % formamide, 5 mM EDTA, 0.02 % bromophenol blue, 0.02 % xylene cyanol), heated at 95°C for 3 min, fractionated on 8% polyacrylamide (19:1)/8.3 M urea gels, and transferred to Amersham Hybond N+ in 0.5x TBE at 400 mA for 1 h. **Northern Blotting.** Northern blotting was performed as previously described (Chen and Wolin, 2023). After transfer, blots were hybridized with [^32^P]-labelled oligonucleotides in Church & Gilbert’s hybridization buffer (0.5 M NaPO_­­­­_pH 6.8, 1 mM EDTA and 7% SDS) at 39°C. Radioactive signals were scanned using a Typhoon FLA 7000 Phosphorimager (GE Healthcare). Quantitation was performed using ImageJ software (NIH). The charged fraction of tRNA was calculated using the equation charged fraction = charged band/(charged band + uncharged band). Oligonucleotide probes for *Msm* tRNAs were: tRNA^Ala(CGC)^: 5′-CTACTCGATGCGAACGAGTCGCGCTAC-3′ tRNA^Ala(GGC)^: 5′-CCCCACACTGCCAGTGTGGTGCGCTAC-3′ tRNA^Arg(TCT)^: 5′-CCTGCAACCGTCGGATTAGAAGGCCGAT-3′ tRNA^His(GTG)^: 5′-GGATCACAACCTGGTGCTCTACCAACTGAACTACAGCCAC-3′ tRNA^Val(CAC)^: 5′-GAGGCGGACGCTCTCCCACTGAGCTACGAGACC-3′ 5S rRNA: 5′-CTTAGCTTCCGGGTTCGGGATGGGACCGGG CGTTTCC-3′

### Flow cytometry

Mid-log phase cultures grown in 7H9 medium were washed once with PBS containing 0.02% tyloxapol, and pellets were resuspended in 500 µL of PBS. Tubes were then centrifugated for three minutes at 400 rpm to pellet bacterial aggregates. 400 µL of the supernatant was transferred to a new tube and diluted with PBS to a final OD of 0.1. Samples were analysed on a BD LSRFortessa x20 cell analyzer (BD Biosciences). mScarlet fluorescence was detected using a yellow-green laser (561 nm). Data were analysed using FlowJo, and graphs were created using R.

### Statistical analysis

Statistical analysis was done with GraphPad Prism 10. Unless otherwise specified, data is presented as mean +/− standard deviation of at least three independent biological replicates per condition. Where appropriate, statistical significance was calculated with either student’s t-test (two-tailed, unpaired) or one-way ANOVA (for multiple groups, one independent variable).

---

**Table S1.** Transcripts differentially regulated in the ups_A_G high survival mutant compared to WT *Msm*.

**Table S2.** Coding sequences for wild-type (WT), GCC_to_GCG and no_ala *whiB7* uORFs. Red: alanine codon that forms part of the anti-terminator loop. Underlined codons and amino acids correspond to the positions of alanine codons and residues in the WT uORF. Codons and amino acids in bold indicate modifications from the native uORF sequence.

**Table S3.** List of primers and strains used in this study.

## ACKNOWLEDGEMENTS

We thank Peter WS Hill (King’s College London); Kristine Arnvig (University College London); José Penadés and D. Holden (Centre for Bacterial Resistance Biology, Imperial College London); Teresa Cortes, Iñaki Comas, and Álvaro Chiner-Oms (Instituto de Biomedicina de Valencia CSIC); Beth Sawyer (University of Westminster), and Julien Hervé for assistance and advice; Carl Nathan (Weill Cornell Medicine) for critical reading of this manuscript; and Nicholas C Poulton and Jeremy Rock (Rockefeller University) for sharing the mScarlet construct and CRISPRi plasmids. We thank the IPBS-TRI, member of the national infrastructure France-BioImaging, which is supported by the French National Research Agency (ANR-10-INBS-04). Funding: This work was supported by the Department of Infectious Disease at Imperial College London (to J.V.), the UK Academy of Medical Sciences under award number SBF006\1172 (to J.V.), the Potts Memorial Foundation (to J.V.), a European Commission Marie Skłodowska-Curie Actions Individual Fellowship (to H.B.), a Medical Scientist Training Program grant from the National Institute of General Medical Sciences of the National Institutes of Health under award number T32GM007739 to the Weill Cornell/Rockefeller/Sloan Kettering Tri-Institutional MD-Ph.D. Program (to S.M.S.), an F30 Predoctoral Fellowship from the National Institute of Allergy and Infectious Diseases of the National Institutes of Health under award number F30AI140623 (to S.M.S.), and a National Defence Science and Engineering Graduate Fellowship from the U.S. Department of Defence (to S.M.S.). The content of this study is solely the responsibility of the authors and does not necessarily represent the official views of the National Institutes of Health. X. C. and S. L. W. are supported by the Center for Cancer Research, National Cancer Institute, National Institutes of Health Intramural Research Program (Project ZIA BC011757). The content of this publication does not necessarily reflect the views or policies of the Department of Health and Human Services, nor does mention of trade names, commercial products or organizations imply endorsement by the US Government. Author contributions: Conceptualization: A.M., H.B., and J.V. Methodology: A.M., H.B., S.M.S., Y.L., X.C., E.J., J.T., P.P. and J.V. Resources: S.W., C.C., P.P., and J.V. Visualization: A.M., H.B. and J.V. Formal analysis: A.M., H.B., P.P., S.W., and J.V. Project administration: J.V. Supervision: H.B. and J.V. Writing—Original draft: H.B. and J.V. Writing—Review and editing: A.M., H.B., S.M.S. and J.V.. Competing interests: The authors declare that they have no competing interests. Data and materials availability: All data needed to evaluate the conclusions in the paper are present in the paper and/or the Supplementary Materials. All bacterial strains listed in this paper can be provided by J.V.’s lab pending scientific review and a completed material transfer agreement. Requests should be directed to J.V. at.

